# Biochemical Upcycling of PET via Glycolysis and Engineered Microbial Consortia

**DOI:** 10.64898/2025.12.04.692332

**Authors:** Francisco J. Molpeceres-García, Alejandro García-Miró, Alicia Prieto, David Sanz, Jorge Barriuso

## Abstract

Polyethylene terephthalate (PET) waste remains a major environmental challenge due to its recalcitrance and low economic value. Here, we present an integrated biochemical approach that couples glycolysis with a synthetic microbial consortium to upcycle post-consumer PET (pcPET) into polyhydroxyalkanoates (PHA). Glycolysis efficiently depolymerized pcPET into bis(2-hydroxyethyl) terephthalate (BHET) in 2 h, circumventing the limitations of *in vivo* PET degradation. We engineered a two-species microbial consortium composed of *Comamonas testosteroni* RW31, able to metabolise terephthalic acid, and *Pseudomonas putida* JM37, able to consume ethylene glycol, each modified for the extracellular secretion of PET- and MHET-hydrolases, employing different plasmid architectures. This division of labour enabled rapid BHET hydrolysis and the subsequent upcycling of the released monomers into PHAs. The combination of the different strains allowed to select *C. testosteroni* pSEVA354-MHETase and *P. putida* pSEVA234-PETase as the best consortium, based on growth and PHAs content. Overall, this work proposes a strategy for PET waste depolymerisation and valorisation, highlighting the potential of mixed chemical and biological approaches and the use of non-conventional microbial chassis within engineered consortia.

## 1. Introduction

Plastics are fundamental for modern societies, but their extensive use, combined with their resistant nature and poor management, causes serious environmental problems (Bucci et al., 2020; Thushari & Senevirathna, 2020). Furthermore, their implications in human health, although not well understood yet, are concerning (Nihart et al., 2025; Wright & Kelly, 2017). There are various causes for its negligent management, from insufficient infrastructure to the lack of economic incentives for recycling: usually, recycled plastic is more expensive, and its properties are worse than virgin plastics, making it less valuable (OECD, 2022). Given this context, the development of strategies for profitable plastic waste utilization has gained interest. Biotechnological approaches to this challenge aim to employ bioprocesses to recycle or upcycle plastic waste, meaning to use it as feedstock to obtain valuable products (Carniel et al., 2024; Ellis et al., 2021). Within this research field, the valorisation of polyethylene terephthalate (PET) has developed the most. PET is a polyester formed by alternating ethylene glycol (EG) and terephthalic acid (TPA) monomers, which is mainly employed for single-use products related to packaging (OECD, 2022). There is a number of described enzymes and microorganisms able to depolymerize this chemical structure (Choi et al., 2023; Yao et al., 2024). However, PET depolymerization efficiency reduces drastically with crystallinity and at reaction temperatures below its glass transition temperature (Tg), ∼60 °C in water (Carniel et al., 2024), which poses a serious bottleneck for *in vivo* degradation. Alternatively, *in vitro* biological depolymerization uses thermophilic enzymes active at temperatures near the PET glass transition temperature (Eugenio et al., 2022; Sonnendecker et al., 2022; Sulaiman et al., 2012). However, the low efficiency of the known enzymes against non-amorphous PET limits their use to degrade post-consumer PET (pcPET) (Amalia et al., 2024). On the other hand, chemical depolymerization, or chemolysis, can easily degrade pcPET through a variety of strategies such as hydrolysis, aminolysis, alcoholysis (Cao et al., 2022), or glycolysis (Feng et al., 2025). These methodologies degrade PET either fully or partially, yielding monomers (EG, TPA), intermediates such as bis-(2-hydroxyethyl)-terephthalate (BHET) or PET oligomers (Barnard et al., 2021). Glycolysis is the most widely applied of these methods, using EG to solubilize and break down PET into BHET. This intermediate facilitates efficient repolymerization to produce high-quality PET (Feng et al., 2025), but these procedures often require high temperatures and pressures or hazardous catalysts, reducing their sustainability. Current research in this field aims to develop milder processes to manage PET waste, using green catalysts to reduce the energy costs and environmental impact of industrial recycling (Barnard et al., 2021).

Within this framework, microorganisms can play an essential role, since several of them can assimilate TPA and/or EG to be transformed into a variety of valuable compounds (Diao et al., 2023a; Johnson et al., 2025; Kim et al., 2019; Molpeceres-García et al., 2025b; Molpeceres-García et al., 2025a; Sadler & Wallace, 2021; Welsing et al., 2025). Thus, the PET monomers can be used as feedstock for biotransformations mediated by a single organism (mono-cultures) or by microbial consortia (Bao et al., 2023). The biotechnological use of consortia offers potential advantages over pure cultures, as their wider genetic repertoire can expand the bioprocess’ possibilities. Some examples are the activation of metabolic pathways not expressed in mono-cultures (Netzker et al., 2018), the increased production of target molecules or enzymes (Hays et al., 2017) or the enhanced robustness against environmental stress (Brenner & Arnold, 2011). However, the main advantage of consortia is the division of labour, through which tasks are divided between the members of the community. Coupled with synthetic biology, the division of labour allows to distribute the burden of expressing heterologous proteins, potentially increasing the efficiency (McCarty & Ledesma-Amaro, 2019). In the case of PET, the availability of two different carbon sources (TPA and EG) eases the design of consortia, as competition for carbon is avoided (McCarty & Ledesma-Amaro, 2019).

In previous studies, we have characterized two bacteria as synthetic biology chassis in the context of PET upcycling: *Comamonas testosteroni* RW31, which can use TPA to accumulate polyhydroxybutyrate (PHB) (Molpeceres-García et al., 2025a), and *Pseudomonas putida* JM37, which can grow on EG and accumulates medium chain length polyhydroxyalkanoates (mcl-PHA) (Molpeceres-García et al., 2025b). In this work, we have designed a microbial consortium in which these bacteria have been modified to secrete the FAST-PETase (Lu et al., 2022) and the *Ideonella sakaiensis* MHETase (*Is*MHETase) (Yoshida et al., 2016), two enzymes that work synergically to degrade PET to TPA and EG. Furthermore, we have employed different plasmid architectures to show their effects on the population dynamics of the consortia. To demonstrate its applicability, we propose a chemo-biological approach for PET upcycling to PHB and mcl-PHA, combining glycolysis of pcPET with fermentation of BHET by a selected consortium.

## 2. Materials and methods

### 2.1. Microorganisms and culture conditions

The bacterial strains and plasmids used in this work are listed in Table 1. Unless indicated, microorganisms were grown in LB medium and transferred to LB or MC minimal medium (KH2PO4 0.33 g/L, Na2HPO4 1.2 g/L, NH4Cl 0.1 g/L, MgSO4*7H2O 0.1 g/L, CaCl2 44 mg/L, adjusted to pH 7). The vitamins and trace elements solutions are described in Barragán et al. (2004). When necessary, 50 µg/mL of kanamycin (Km) and 34 µg/mL of chloramphenicol (Cr) were added to the growth media. Cultures were grown at 30 °C and 200 rpm and gene expression was induced with 1 mM IPTG.

**Table 1.**
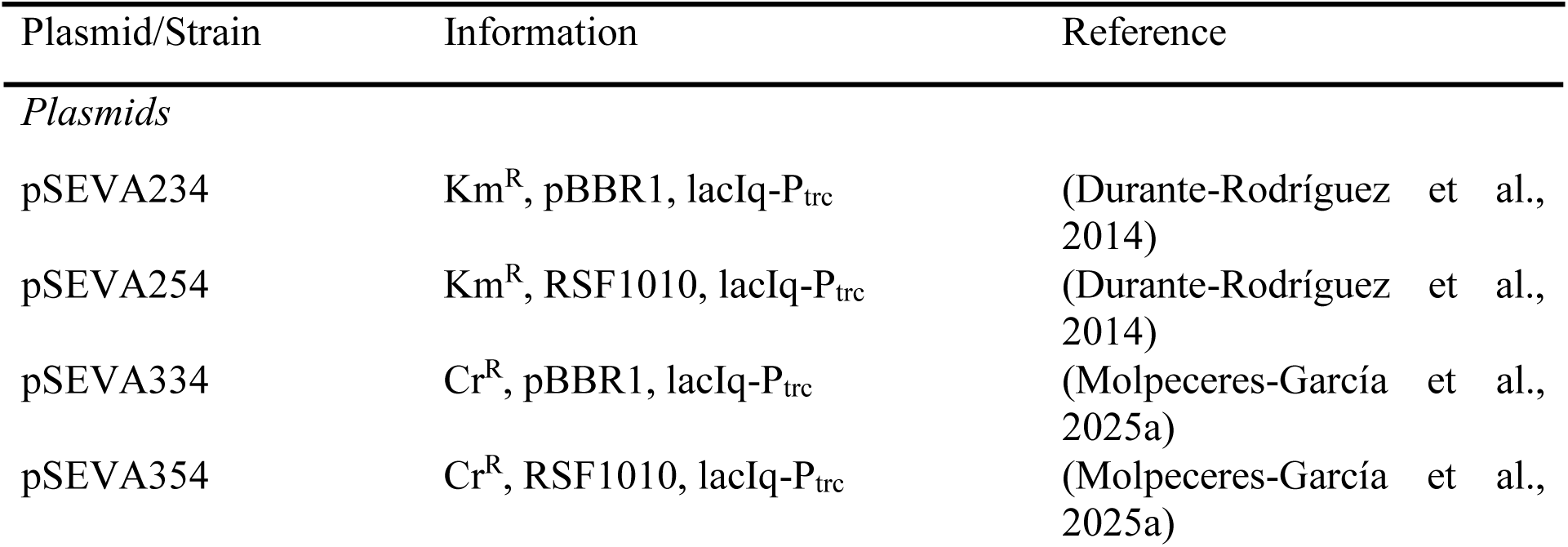
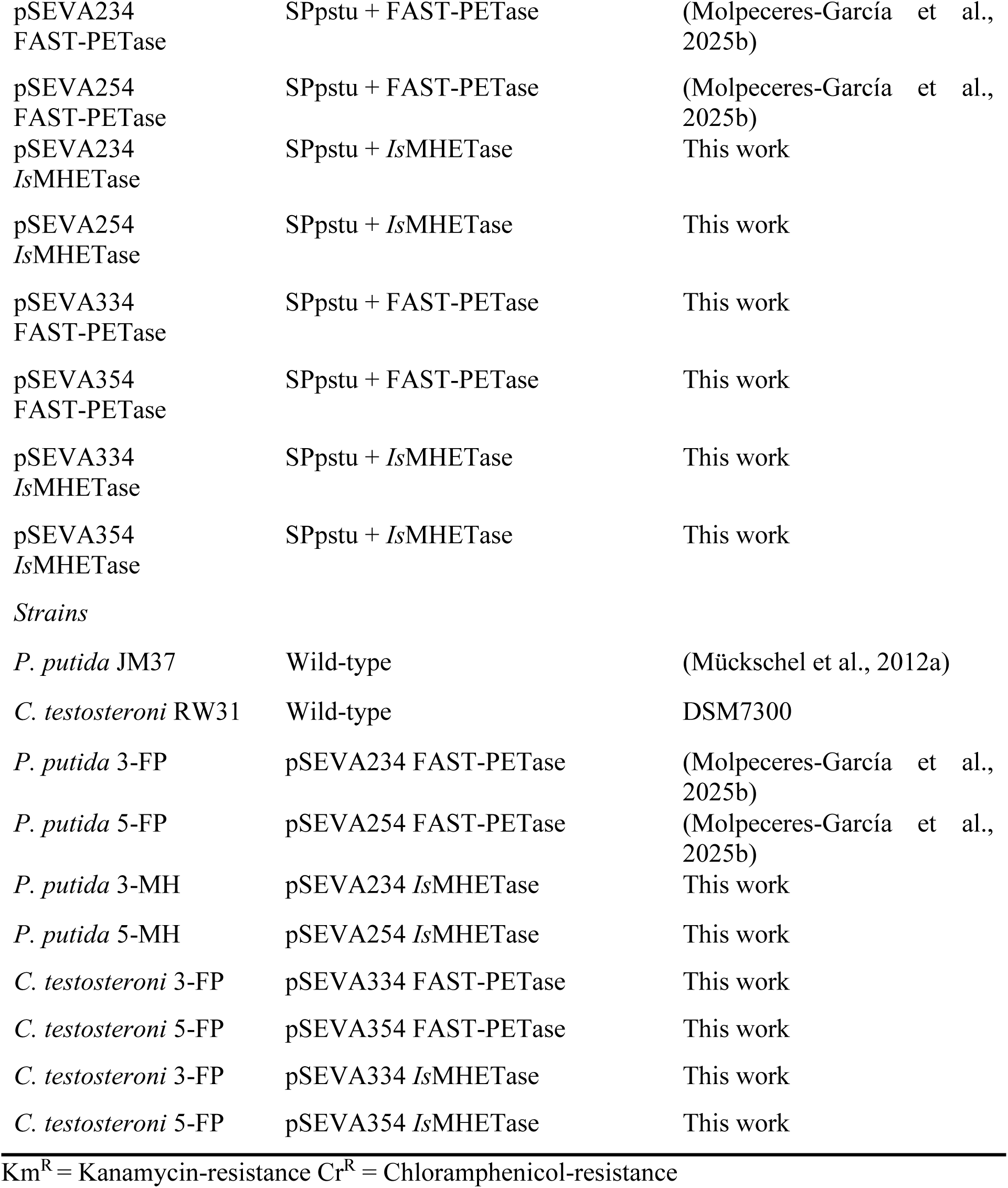
Strains and plasmids employed in this study

| Plasmid/Strain | Information | Reference |
| --- | --- | --- |
| *Plasmids* |  |  |
| pSEVA234 | Km<sup>R</sup>, pBBR1, lacIq-P<sub>trc</sub> | (Durante-Rodríguez et al., 2014) |
| pSEVA254 | Km<sup>R</sup>, RSF1010, lacIq-P<sub>trc</sub> | (Durante-Rodríguez et al., 2014) |
| pSEVA334 | Cr<sup>R</sup>, pBBR1, lacIq-P<sub>trc</sub> | (Molpeceres-García et al., 2025a) |
| pSEVA354 | Cr<sup>R</sup>, RSF1010, lacIq-P<sub>trc</sub> | (Molpeceres-García et al., 2025a) |
| pSEVA234<br>FAST-PETase | SPpstu + FAST-PETase | (Molpeceres-García et al., 2025b) |
| pSEVA254<br>FAST-PETase | SPpstu + FAST-PETase | (Molpeceres-García et al., 2025b) |
| pSEVA234<br>*IsMHETase* | SPpstu + *IsMHETase* | This work |
| pSEVA254<br>*IsMHETase* | SPpstu + *IsMHETase* | This work |
| pSEVA334<br>FAST-PETase | SPpstu + FAST-PETase | This work |
| pSEVA354<br>FAST-PETase | SPpstu + FAST-PETase | This work |
| pSEVA334<br>*IsMHETase* | SPpstu + *IsMHETase* | This work |
| pSEVA354<br>*IsMHETase* | SPpstu + *IsMHETase* | This work |
| *Strains* |  |  |
| *P. putida* JM37 | Wild-type | (Mückschel et al., 2012a) |
| *C. testosteroni* RW31 | Wild-type | DSM7300 |
| *P. putida* 3-FP | pSEVA234 FAST-PETase | (Molpeceres-García et al., 2025b) |
| *P. putida* 5-FP | pSEVA254 FAST-PETase | (Molpeceres-García et al., 2025b) |
| *P. putida* 3-MH | pSEVA234 *IsMHETase* | This work |
| *P. putida* 5-MH | pSEVA254 *IsMHETase* | This work |
| *C. testosteroni* 3-FP | pSEVA334 FAST-PETase | This work |
| *C. testosteroni* 5-FP | pSEVA354 FAST-PETase | This work |
| *C. testosteroni* 3-FP | pSEVA334 *IsMHETase* | This work |
| *C. testosteroni* 5-FP | pSEVA354 *IsMHETase* | This work |

Km<sup>R</sup> = Kanamycin-resistance Cr<sup>R</sup> = Chloramphenicol-resistance

### 2.2. TPA and EG tolerance

*C. testosteroni* RW31 and *P. putida* JM37 were grown in MC minimal medium with 10 mM TPA (Sigma-Aldrich, Ref 185361) or EG (Sigma-Aldrich, Ref 102466), respectively, as carbon sources. To assess *C. testosteroni* RW31 tolerance to EG, the bacterium was grown with 0 mM, 10 mM, 25 mM and 50 mM EG. In the case of *P. putida* JM37, TPA was tested at the same concentrations. *C. testosteroni* RW31 tolerance to TPA and *P. putida* JM37 tolerance to EG have been evaluated in previous works (Molpeceres-García et al., 2025b; Molpeceres-García et al., 2025a). Growth curves were performed in a 96 well plate using a Victor Nivo Multimode Plate Reader (PerkinElmer) at 30 °C and 200 rpm. Growth was measured as optical density at 600 nm (OD^600^). Bacteria were inoculated at 0.1 OD^600^.

### 2.3. Plasmid design and transformation

Codon-optimized FAST-PETase and *Is*MHETase with SPpstu signal peptide from *Pseudomonas stutzeri* MO-19 (Fujita et al., 1989) were cloned into pSEVA234, pSEVA254, pSEVA334 and pSEVA354 plasmids, using *Eco*RI and *Hind*III restriction sites (Durante-Rodríguez et al., 2014). *P. putida* JM37 was transformed with Km^R^ plasmids, whereas *C. testosteroni* RW31 was transformed with Cr^R^ plasmids. The resulting strains were named *P. putida* 3-FP, *P. putida* 5-FP, *P. putida* 3-MH, *P. putida* 5-MH, *C. testosteroni* 3-FP, *C. testosteroni* 5-FP, *C. testosteroni* 3-MH and *C. testosteroni* 5-MH (Table 1).

Plasmids were transformed by electroporation. First, electrocompetent cells were prepared as explained in Molpeceres-García et al. (2025b). Electroporation was performed in 0.2 cm cuvettes with a MicroPulse elctroporator (BioRad) (2.5 kV, 200 Ω, 25 µF) by adding 100 ng of plasmid. Cells were recovered in LB for 2 h and plated in LB with either 50 µg/mL kanamycin or 34 µg/mL chloramphenicol for selection.

### 2.4. PETase activity assay

The FAST-PETase activity in *P. putida* 3-FP and 5-FP strains was reported in a previous work (Molpeceres-García et al., 2025b). To assess *C. testosteroni* 3-FP and 5-FP strains activity, they were cultured overnight in LB and then inoculated at 0.1 OD^600^ in fresh LB with 1 mM IPTG. Chloramphenicol was used for plasmid maintenance. The non-induced recombinant strains and the wild type strains were used as negative controls. After 48 h, the extracellular esterase activity was measured colorimetrically, following the release of *p*-nitrophenol from 1.5 mM *p*-nitrophenyl-butyrate (*p*NPB) (Sigma-Aldrich, Ref N9876) through the increase in absorbance at 410 nm. Reactions were conducted in 50 mM phosphate buffer pH 7.5 using 100 µL of culture supernatant. One unit of activity is defined as the amount of enzyme used to release 1 μmol of *p*-nitrophenol (ε410 =15,200 M^-1^ cm^-1^) per minute.

### 2.5. MHETase activity assay

As MHETase activity cannot be determined following *p*NP-aliphatic esters hydrolysis (Yoshida et al., 2016), *in vivo* assays were performed. *P. putida* 5-FP was co-inoculated with either *P. putida* 3-MH or *P. putida* 5-MH, each at 0.05 OD^600^, in MC minimal medium with 0.5 g/L yeast extract as nitrogen source and 10 mM EG as carbon source, adding 5 mM of BHET (Sigma-Aldrich, CAS 959-26-2) and 1 mM IPTG. A similar assay was performed for co-cultures of *C. testosteroni* 5-FP with 3-MH or 5-MH, using 10 mM TPA as carbon source. Monocultures of each strain were also analyzed as negative controls. The resulting monomers were measured by HPLC.

### 2.6. PET monomers quantification by HPLC

An Agilent 1200 series HPLC was used for EG, BHET, MHET and TPA quantification. EG concentration was measured using an RID detector (Agilent), a BioRad Aminex HPX-87H column (300 x 7.8 mm), and H2SO4 2 mM as the mobile phase, under isocratic conditions at 0.5 mL/min over 25 min. The column was maintained at 60 °C and RID at 35 °C. EG retention time was 19.5 min. For BHET, MHET and TPA analysis, samples were first diluted 1:1 with acetonitrile, to ensure proper dissolution of BHET and MHET. Then, they were separated on a ZORBAX Eclipse Plus C18 column (4.6 × 250 mm) and detected at 260 nm using a DAD detector. The solvents used for the mobile phase consisted of Milli Q water with 0.1 % acetic acid (A) and acetonitrile with 0.1 % acetic acid (B). The column was initially equilibrated with 95 % A and 5 % B. Elution was made with a gradient from 95 % to 30 % A over 25 min. The flow rate was maintained at 0.8 mL/min and the column temperature at 30 °C. Retention times were 10.2 min for TPA, 11.4 min for MHET and 12.1 min for BHET.

### 2.7. Post-consumer PET glycolysis

Glycolysis was carried out at 200 °C in a 250 mL flask containing 75 mL of EG and 6 g of post-consumer PET (pcPET) obtained from a commercial water bottle (Nestlé Aquarel®). Zn(NO3)2 was added as catalyst (1% w/v). pcPET was cut into ∼1x1 cm pieces and added to the flask once the EG reached the proper temperature. The reaction was carried out for 2.5 h, sampling every 30 min to monitor BHET formation, and allowed to cool before recovery of the product (post-consumer BHET, pcBHET) by crystallization. To do so, Milli-Q water was added in 1:10 proportion and the solution stored at 4 °C overnight. pcBHET crystals deposited at the bottom of the flask were recovered after filtering through filter paper and washing with Milli-Q water, and dried at 50 °C. Finally, the crystals were ground with a mortar and passed through a 0.42 mm sieve. The pcBHET molar yield of the reaction was calculated according to the equation

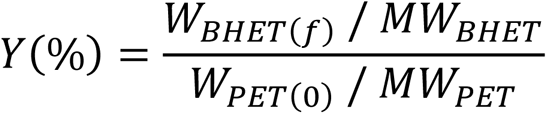

where WBHET(f) refers to the final recovered BHET weight, WPET(0) refers to the initial PET weight and MW refers to the molecular weight (López-Fonseca et al., 2010).

### 2.8. Consortia cultivation and growth monitoring

Experiments were carried out in 250 mL flasks filled with 50 mL of MC medium supplemented with 5 mM TPA and 5 mM EG. Each strain was inoculated at 0.05 OD^600^ and the cultures incubated for 48 h at 30 °C and 180 rpm. The consortia used are listed in Table 2. Either 5 mM of BHET or pcBHET was used to test the conversion efficiency consortia. Samples were taken at 0, 6, 24 and 48 h for PET monomer quantification, PHAs content analysis and growth assessment. To monitor *C. testosteroni* RW31 and *P. putida* JM37 growth separately the plate counting method was used. For that, three 10 µL drops were plated on Petri dishes containing MC-agar supplemented with either 10 mM TPA or 10 mM EG. Plates were incubated 24 h at 30 °C and then growth was measured as colony forming units per mL (CFU/mL).

**Table 2.** Consortia employed in this work.

| Consortia | *C. testosteroni* strain | *P. putida* strain |
| --- | --- | --- |
| Con1 | 3-MH | 3-FP |
| Con2 | 3-FP | 3-MH |
| Con3 | 5-MH | 5-FP |
| Con4 | 5-FP | 5-MH |
| Con5 | 3-MH | 5-FP |
| Con6 | 5-MH | 3-FP |
| Con7 | 5-FP | 3-MH |
| Con8 | 3-FP | 5-MH |

### 2.9. PHA analysis and quantification by GC-MS

PHA production was visually confirmed by Nile Red staining (Zuriani et al., 2013). Briefly, 500 µL of culture were centrifuged at 7,000 rpm, resuspending the resulting pellet in 50 µL of PBS and adding Nile Red to a final concentration of 5 µg/mL. PHA granules were observed with a DM4 B microscope (Leica) equipped with a *p*E-300 Lite LED illuminator (CoolLED) and a DFC345 FX camera (Leica). The consortium in which more PHA content was observed was selected for monomer identification and quantification. To analyse the composition and quantify total PHA, 2-5 mg of freeze-dried biomass were methanolized. For each biological sample, two technical replicates were included. Samples were submitted to two different treatments, in order to properly extract PHB and mcl-PHA. Briefly, freeze-dried pellets were resuspended in 2 mL of methanol containing 3% of sulfuric acid (15% for mcl-PHA). Then, 2 mL of 3-methylbenzoic acid, dissolved in chloroform at 0.5 mg/mL, were added as an internal standard. Samples were incubated at 100 °C for 4 h (5 h for mcl-PHA). Reactions were stopped by cooling in ice. Samples were washed twice by adding 2 mL of Milli-Q water, centrifuging for phase separation and discarding the aqueous phase and the cellular debris. For mcl-PHA samples, a third washing step was done with a saturated solution of Na2CO3 to eliminate residual sulfuric acid. Finally, a small amount of Na2SO4 was added to completely remove water content. Methanolized samples were analysed with an Agilent (Santa Clara, California, USA) INTUVO 9000 GC System gas chromatograph coupled with a 5977C GC/MSD detector (EI, 70 eV). During this work, the HP-5MS UI column (30 m × 0.25 mm x 0.25 μm film thickness) was used. For PHB, the starting temperature was 60 °C, with an increase of 5 °C/min until 65 °C and 15 °C/min until 125 °C, maintaining this temperature for 3.5 min. The retention time for the methyl ester of the C4 monomer was 2.4 min, and for the internal standard 7.7 min. For mcl-PHA, the starting temperature was 80 °C and it was increased 15 °C/min up to 175 °C, holding 20 min at this temperature. Retention times for the different methyl ester monomers were: 3.5 min (C6), 5.2 min (C8), 7.0 min (C10), 9.1 min (C12:1) and 9.4 min (C12). Internal standard was detected at 4.9 min.

## 3. Results and discussion

### 3.1. C. testosteroni RW31 and P. putida JM37 are suitable for a consortia

In order to develop a PET-upcycling consortium, we first assessed TPA and EG toxicity in *P. putida* JM37 and *C. testosteroni* RW31, respectively. With 50 mM EG, *C. testosteroni* showed a delay of 18 h to reach the stationary phase and a decrease of ∼7 % in its maximum OD^600^, whereas with 10 and 25 mM almost no differences, compared to the positive control, were detected (Fig. 1A). In the case of *P. putida* JM37, the effect of TPA was more marked, with a decrease in the maximum OD^600^ of ∼18 % with 10 and 25 mM of TPA and ∼22 % with 50 mM (Fig. 1A). Hence, both microorganisms were able to tolerate TPA and EG, although *P. putida* JM37 seems to be more sensitive to TPA. Growth inhibition by TPA has been observed in a modified strain of *P. putida* KT2440 able to assimilate EG, in which increasing concentrations of TPA led to a decrease in EG consumption and bacterial growth (Bao et al., 2023). The same study demonstrated that the formation of a consortium with a second *P. putida* population, modified to assimilate TPA, relieved its inhibitory action. In our case, the co-culture of *P. putida* JM37 with the TPA-consumer *C. testosteroni* RW31 may have a similar effect.

**Figure 1.**
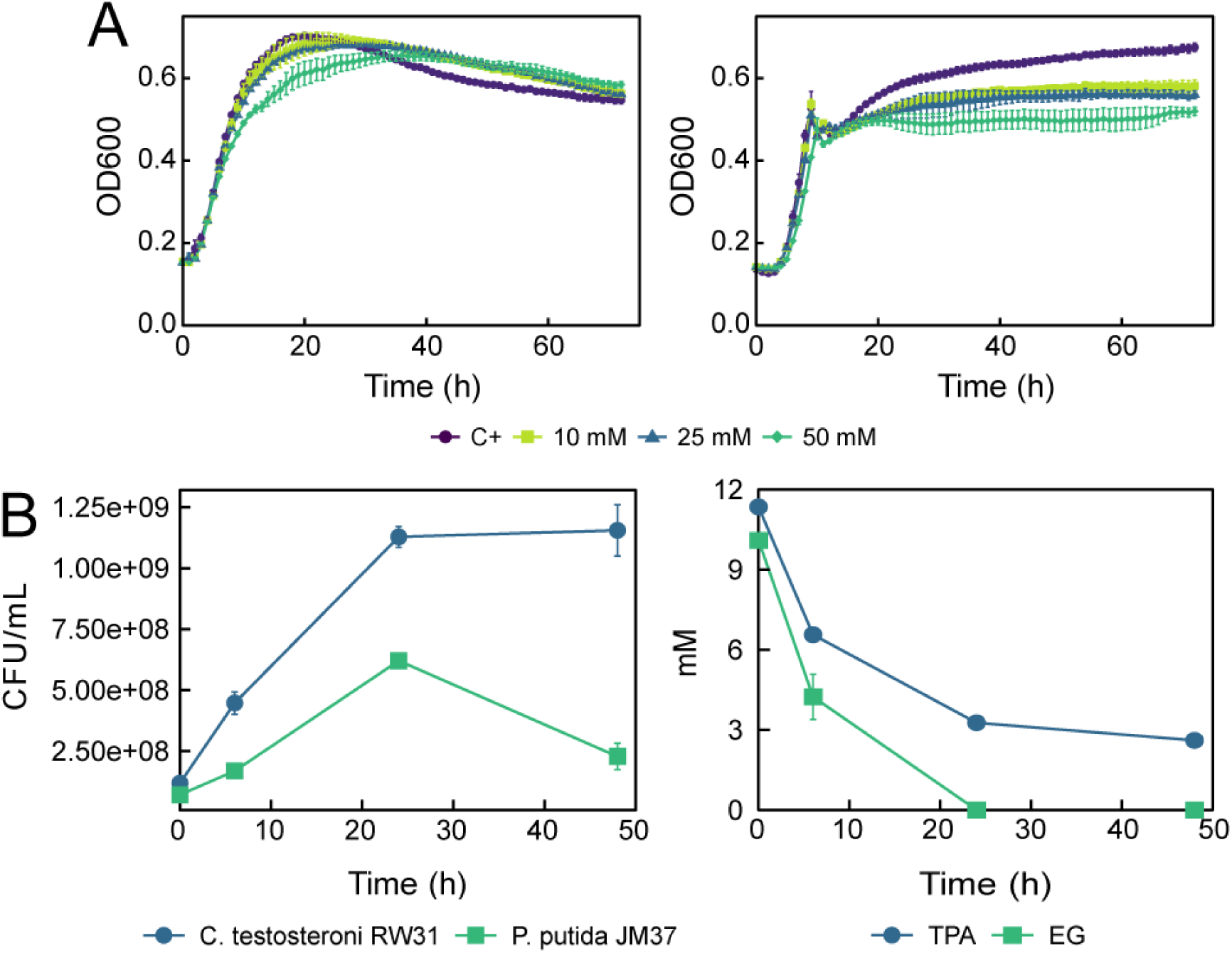
Growth of the wild-type strains *C. testosteroni* RW31 and *P. putida* JM37. (A) Effect of EG on *C. testosteroni* RW31 growth in MC minimal medium with 10 mM TPA as carbon source (left); effect of TPA on *P. putida* JM37 growth in MC minimal medium with 10 mM EG as carbon source (right). The legend indicates the concentration of EG or TPA added to the media. (B) Growth of *C. testosteroni* RW31 and *P. putida* JM37 in co-culture in MC minimal medium with 10 mM EG and 10 mM TPA as carbon sources, expressed in CFU/mL (left); TPA and EG consumption in this consortium (right). Results are given as the mean of n = 3. Error bars represent standard error.

Furthermore, *C. testosteroni* RW31 and *P. putida* JM37 demonstrated to grow together in MC minimal medium with 10 mM of TPA and EG as carbon sources. This consortium achieved a cellular density of 1.75x10^9^ CFU/mL in 24 h, composed by a 64.5 % of *C. testosteroni* and a 35.4 % of *P. putida* (Fig. 1B). The prevalence of *C. testosteroni* over *P. putida* could be explained by the imbalance between carbon sources. In this experiment, *P. putida* fully consumed the EG from the culture medium in 24 h, whereas at this time *C. testosteroni* assimilated ∼71% of the available TPA (Fig. 1B), suggesting a carbon excess in TPA and deficit in EG. Moreover, the metabolism of TPA and EG may also play a role in this carbon imbalance. In previous works, we showed that *C. testosteroni* RW31 likely assimilates TPA through protocatechuic acid by the 4,5-*meta* pathway, which yields pyruvate and oxaloacetate, meaning that 7 carbons would be available for each TPA molecule assimilated (Molpeceres-García et al., 2025a; Wilkes et al., 2023). However, *P. putida* JM37 assimilates EG through the Gcl pathway, which yields only one pyruvate per two EG molecules, meaning that each EG would provide 1.5 carbons (Molpeceres-García et al., 2025b; Mückschel et al., 2012). Thus, *C. testosteroni* RW31 would have 4.67 more carbons available than *P. putida* JM37. This suggests that the *C. testosteroni* – *P. putida* proportion could be balanced by adding more EG or a co-substrate to the media. However, we decided to use equimolar concentrations of TPA and EG considering the use of hydrolysated PET as substrate in future experiments, which would yield TPA and EG in 1:1 proportion.

### 3.2. Expression of PET-hydrolases in C. testosteroni RW31 and P. putida JM37

Following our biological approach, we developed a BHET-degrading consortium cloning the FAST-PETase and the *Is*MHETase into *C. testosteroni* RW31 and *P. putida* JM37. Both enzymes were cloned in plasmids with pBBR1 or RSF1010 replication origins to assess whether one resulted better for the protein expression and the consortium fitness than the other, as they differ in copy number (Cook et al., 2018) (Table 1). *C. testosteroni* 3-FP and 5-FP strains presented an extracellular esterase activity against *p*NPB of 0.12 ± 0.007 and 0.28 ±0.007 U/mL, respectively (Fig. 2). *P. putida* 3-FP and 5-FP strains’ activities were reported in a previous work (Molpeceres-García, et al., 2025b), resulting in 0.36 and 0.41 U/mL against *p*NPB, respectively. In both bacteria, RSF1010 plasmids resulted in an improved esterase activity, probably due to its higher copy number. The activity was 3-fold higher in *P. putida* 3-FP compared to *C. testosteroni* 3-FP and 1.46-fold higher in *P. putida* 5-FP compared to *C. testosteroni* 5-FP.

**Figure 2.**
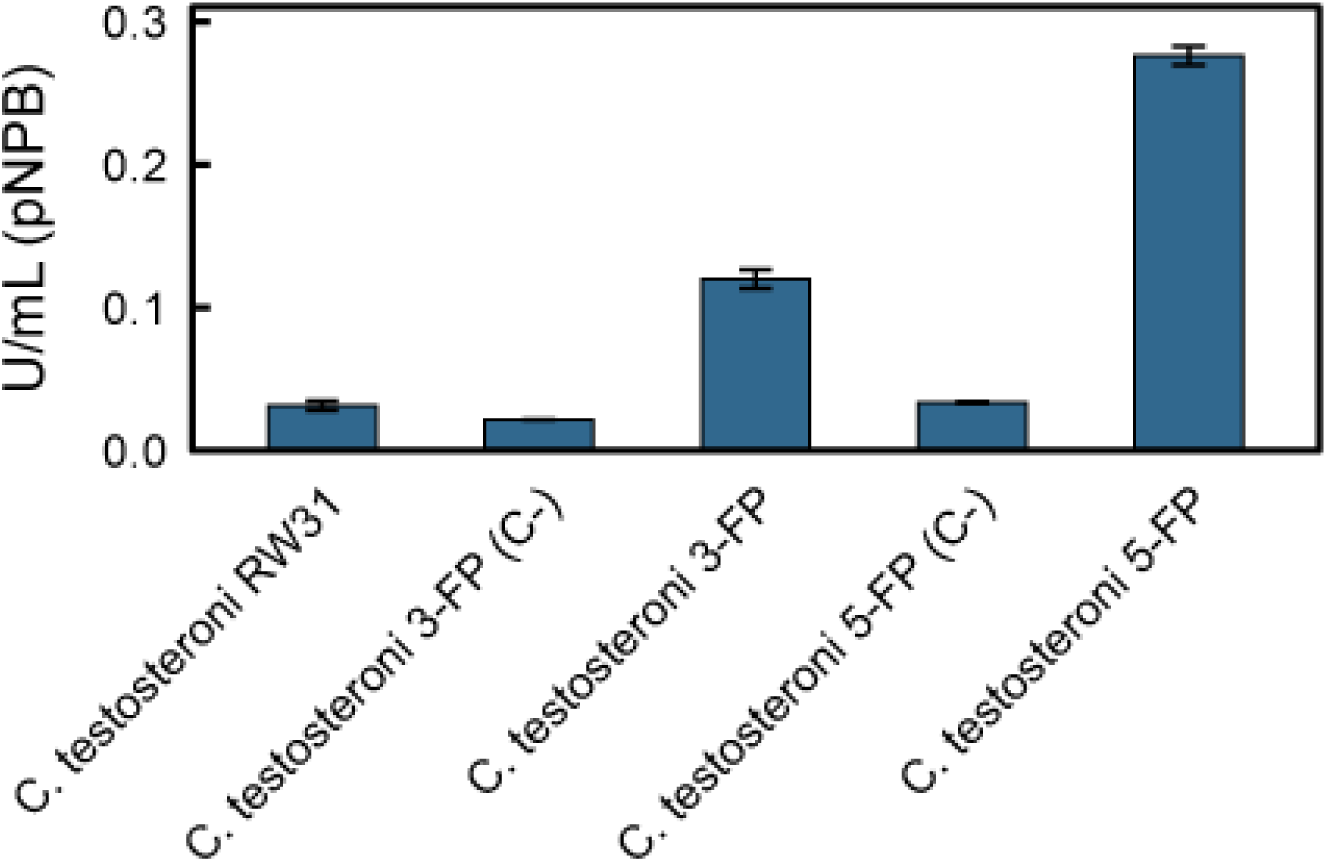
Activity against *p*NPB of supernatants from cultures of *C. testosteroni* 3-FP and 5-FP grown in LB with chloramphenicol for 48 h. C- corresponds to non-induced cultures. Results are given as the mean of n = 2. Error bars represent standard error.

In the case of the MHETase-carrying strains, enzymatic activity was tested indirectly in consortia with a PETase-carrying strain to measure the joint degradation of BHET. Co-cultures of *C. testosteroni* 5-FP with *C. testosteroni* 3-MH or *C. testosteroni* 5-MH degraded 41.2 % and 44.4 % of BHET into EG and TPA, respectively, in 6 h, and the totality in 24 h. MHET was barely detectable during the experiment, indicating that it was quickly degraded. In the case of *P. putida*, the strain 5-FP in consortia with *P. putida* 3-MH or 5-MH degraded BHET completely in 6 h and MHET in 24 h (Fig. 3).

**Figure 3.**
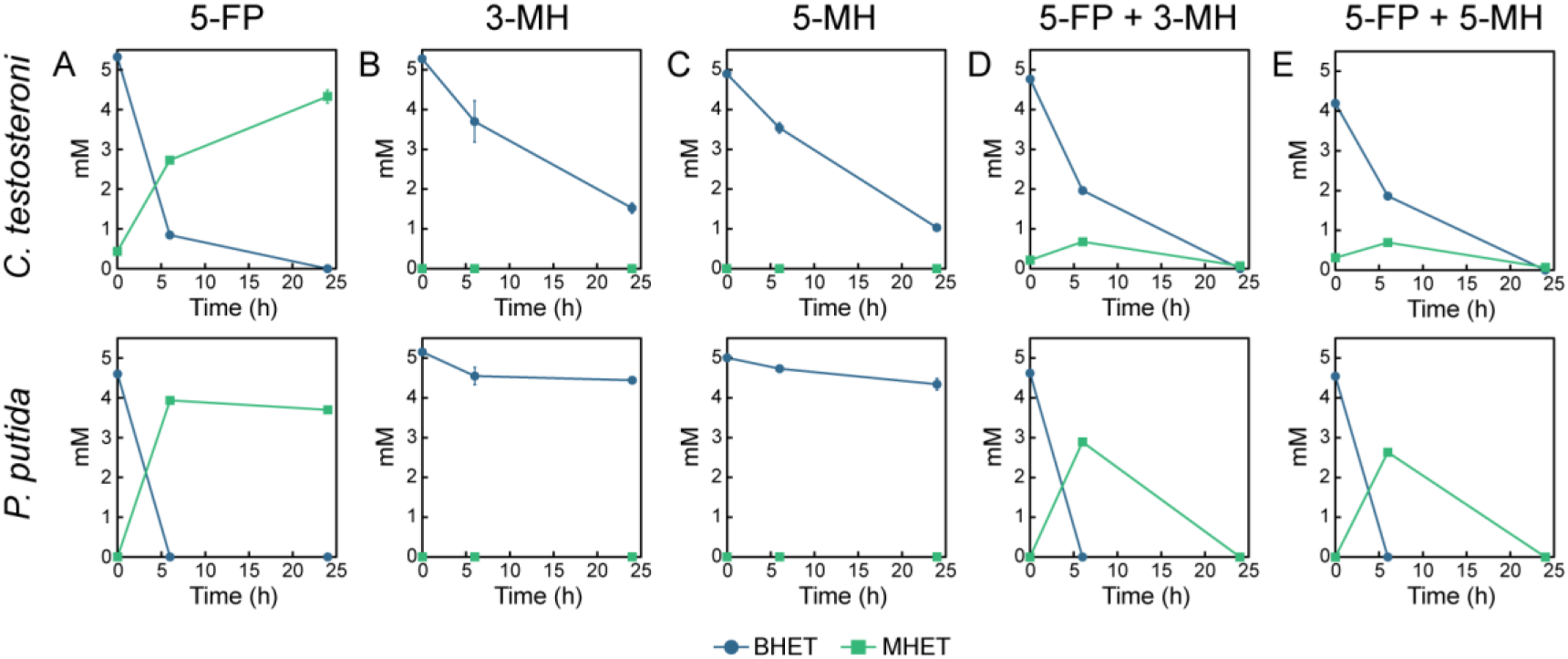
Activity of *C. testosteroni* (up) and *P. putida* (down) recombinant strains against BHET. (A) Mono-culture of 5-FP. (B) Mono-culture of 3-MH. (C) Mono-culture of 5-MH. (D) Co-culture of 5-FP and 3MH. (E) Co-culture of 5-FP and 5-MH. Results are given as the mean of n = 3. Error bars represent standard error.

All the monocultures of MHETase-producing strains induced partial BHET degradation (Fig. 3B, C) including *Is*MHETase, whose activity against BHET is very low (Palm et al., 2019). Since the wild-type *C. testosteroni* RW31 has some degradative effect on BHET (data not shown), this result suggests that, rather than a non-specific PETase activity of *Is*MHETase, this bacterium may secrete an enzyme with low activity against BHET. The faster BHET degradation by *P. putida* 5-FP cultures compared to *C. testosteroni* 5-FP could be explained by its higher secretion of PETase activity. Regarding MHET, all the MHETase-carrying strains showed to have this activity.

### 3.3. Recombinant C. testosteroni RW31 and P. putida JM37 strains can degrade BHET in consortium

Once demonstrated the activity of all the recombinant strains, mixed consortia combining the two species were evaluated for their capacity to grow in MC minimal medium containing BHET as substrate. The previous results show that both *C. testosteroni* RW31 and *P. putida* JM37 can produce PHA in this medium from TPA and EG, respectively. Hence, microbial growth (CFU/mL, Fig. 4) and accumulation of PHA (Nile Red staining, Fig. S1) were evaluated in the consortia listed in Table 2. In all cases, growth was higher for *C. testosteroni* than for *P. putida*, which is consistent with the results obtained with the wild-type consortium (Fig. 1). However, expressing heterologous enzymes did have an effect in the bacterial fitness. While proportions were more equilibrated in Con3, Con5 and Con7, transformation with pSEVA254-*Is*MHETase (Con4, Con8) produced a strong detrimental effect in *P. putida* growth (Fig. 4). This may be explained by the higher copy number of RSF1010 replication origin and the larger size of *Is*MHETase (65 kDa) compared to FAST-PETase (29 kDa) (Palm et al., 2019; Seo et al., 2019), which may represent a high metabolic burden for *P. putida* JM37. Nevertheless, these results suggest that plasmid architectures strongly influence the growth dynamics of *C. testosteroni* RW31 and *P. putida* JM37 recombinant strains. These differences could be useful for a better design of future consortia, as populations’ proportion has been proven important to obtain better performance in synthetic consortia (Bao et al., 2023; Mehta et al., 2025; Qi et al., 2021).

**Figure 4.**
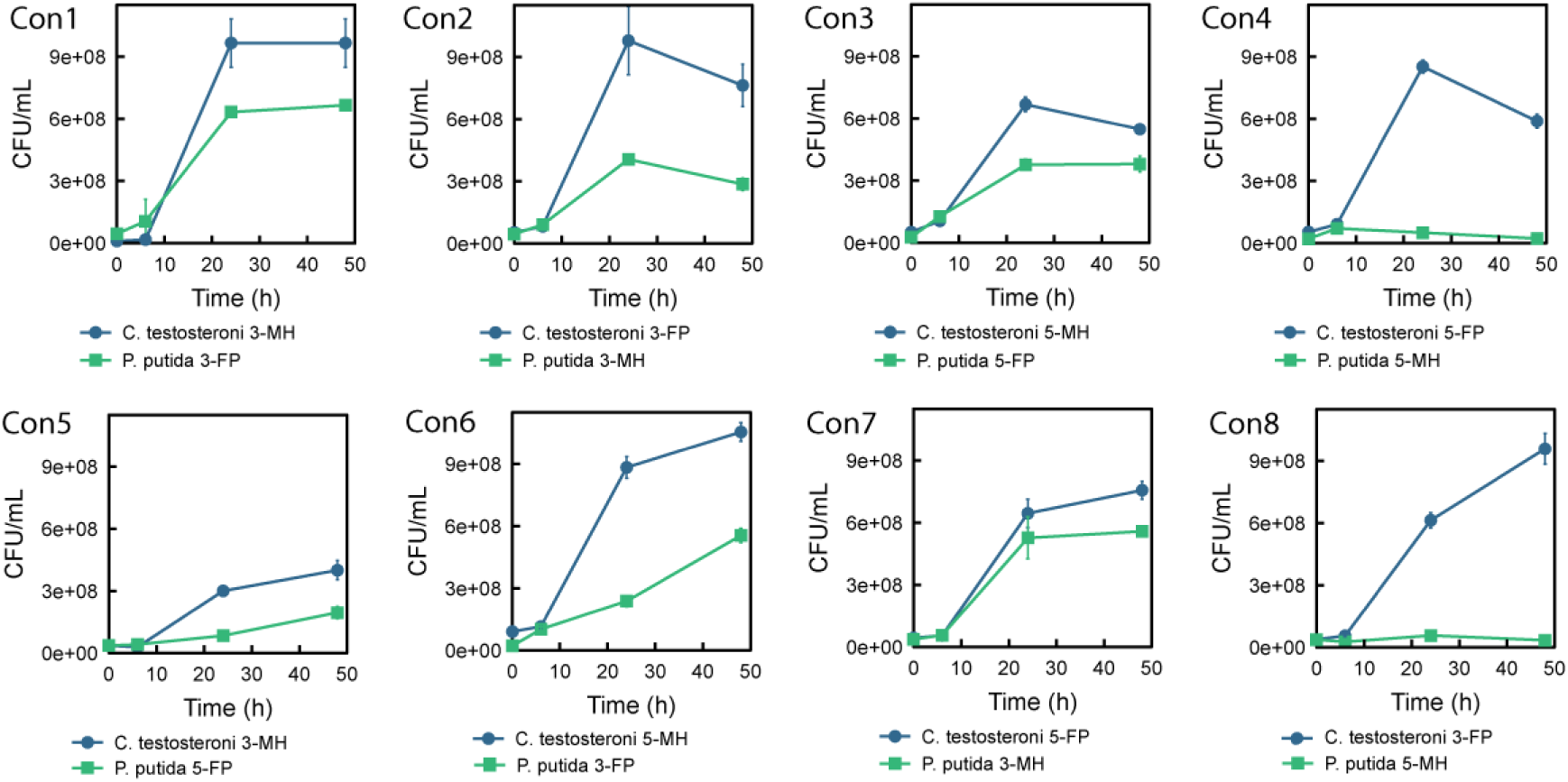
Growth in co-cultures of *C. testosteroni* – *P. putida* recombinant strains in MC minimal media with 5 mM TPA, EG and BHET, expressed as CFU/mL. The strains in each consortium can be consulted in Table 2. Results are given as the mean of n = 3. Error bars represent standard error.

To conduct the following experiments, Con6 (*C. testosteroni* 5-MH + *P. putida* 3-FP) was selected for its high growth (1.12x10^9^ CFU/mL, composed of 78.8% of *C. testosteroni* 5-MH and 21.2% of *P. putida* 3-FP) and PHA accumulation at 24 h (Fig. S1).

Con6 completely degraded 5 mM commercial BHET in 24 h, while MHET was barely present during the experiment (Fig. 5A), demonstrating an effective expression of both FAST-PETase and *Is*MHETase. While EG was fully consumed, only 59.5% of the available TPA was assimilated in 48 h. The populations’ proportion and the TPA/EG consumption of Con6 was like that of the WT strains (Fig. 1), although slower, as it peaked 24 h later. Despite there was 15 mM EG potentially available (5 mM in the media + 10 mM from the BHET hydrolysis), and all of it was consumed, the proportion of *P. putida* was not larger than in the WT consortium. These growth differences could be explained by the metabolic burden of expressing the FAST-PETase. Moreover, in our previous work expressing the FAST-PETase and the MHETase in *C. testosteroni* RW31, we observed that increasing the amount of BHET from 5 to 10 mM in the media impeded bacterial growth, suggesting an inhibitory effect (Molpeceres-García et al., 2025).

**Figure 5.**
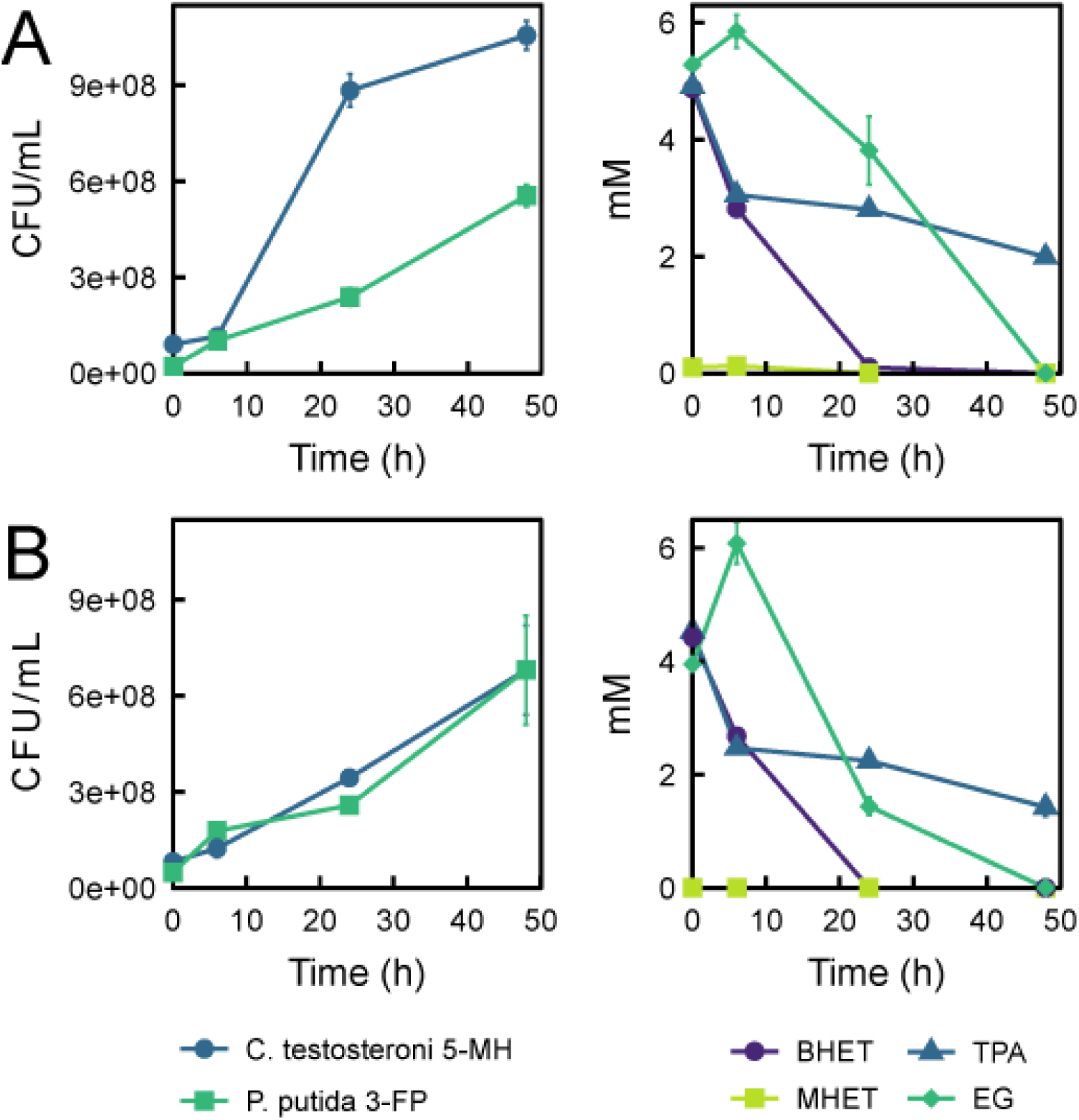
Con6 (*C. testosteroni* 5-MH and *P. putida* 3-FP) growing in MC minimal media containing 5 mM of TPA, EG and (A) BHET or (B) pcBHET. Left panels: growth (CFU/mL). Right panels: BHET, MHET, TPA and EG consumption (mM). Results are given as the mean of n = 3. Error bars represent standard error.

*In vivo* degradation of PET or BHET by microbial consortia has mainly been studied from a top-down perspective. In most reports, workflows involve the isolation of microbial communities from contaminated areas, followed by its characterization and assessment of enzymatic activities and plastic degradation (Gao & Sun, 2021; Maheswaran et al., 2023; Meyer-Cifuentes et al., 2020; Roberts et al., 2020). Alternatively, the bottom-up approach, in which a synthetic consortium is designed for an specific objective, allows to choose suitable organisms with desirable capacities, as well as their enhancement by synthetic biology tools (Jia et al., 2016; Stephens & Bentley, 2020). Qi et al. (2021) designed a multi-species consortium in which *Bacillus subtilis* 168 was modified for expressing either *Is*PETase or *Is*MHETase. In this case, TPA was inhibitory for *B. subtilis*, so *Rhodococcus jostii* RHA1 was added for detoxification. This consortium degraded 7.9 mM of BHET dissolved in a 50% γ-cyclodextrin solution in 20 h and ∼10.5 mg of an amorphous PET film weight in 7 days, growing in LB medium. Also, the addition of *P. putida* KT2440 allowed the assimilation of the released EG. Moreover, the authors reported how the inoculum ratio and medium composition played an important role. This work illustrates the versatility of artificial consortia, but also the complexity of adjusting the cultivation parameters. In our case, the use of TPA- and EG-assimilating organisms as chassis makes the process simpler, as substrate inhibitory effects are not a concern. Furthermore, the fact that each microorganism produces a single degrading enzyme enables cooperation between the consortium members. While *P. putida* 3-FP facilitates the degradation of BHET to MHET, *C. testosteroni* 5-MH degrades MHET, obtaining TPA for its own consumption, providing an extra EG molecule to *P. putida* and eliminating the MHET, which is known to be a PETase inhibitor (Zhang et al., 2023).

### 3.4. Post-consumer PET glycolysis

To obtain pcBHET, 6 g of pcPET were glycolyzed to 940 mM of pcBHET in 2 h (Fig. 6A), with no presence of MHET and TPA in the glycolysis solution. pcBHET was crystallized by adding water to the reaction mixture, obtaining 4.7 g of a solid fraction (59.2 % pcBHET molar yield) and a diluted EG solution. It is worth to mention that an undetermined fraction of the employed pcPET weight can correspond to additives and glues, which could interfere the growth of the microorganisms. This conversion yield could be improved by optimization of parameters such as reaction time, PET-EG molar ratio and the type and concentration of catalyst added (López-Fonseca et al., 2010), or the addition of a cosolvents (Huang et al., 2023). Although residual EG is obtained in the process, it can be purified by distillation and reused for subsequent PET glycolysis. Aguado et al. (2023) performed glycolysis of 200 g PET, reusing the EG up to 10 times, observing a decrease of 6.6 - 7.4 % of pcBHET yield in the last reaction respect to the fresh EG. As an alternative for EG reutilization, we tested weather the residual EG can serve as feedstock to produce PHA in pure cultures of *P. putida* JM37. Adding a 1:10 dilution of the residual EG to MC minimal medium, JM37 grew up to 1.67 ± 0.03 OD^600^ in 48 h and accumulated PHA, as confirmed by Nile Red staining (Fig. 6B, C). Moreover, in a previous work this strain was shown to use EG as carbon source to produce violacein, a highly valuable compound (Molpeceres-García et al., 2025b). Hence, the utilization of the EG recovered after glycolysis can be part of a biochemical strategy for PET upcycling.

**Figure 6.**
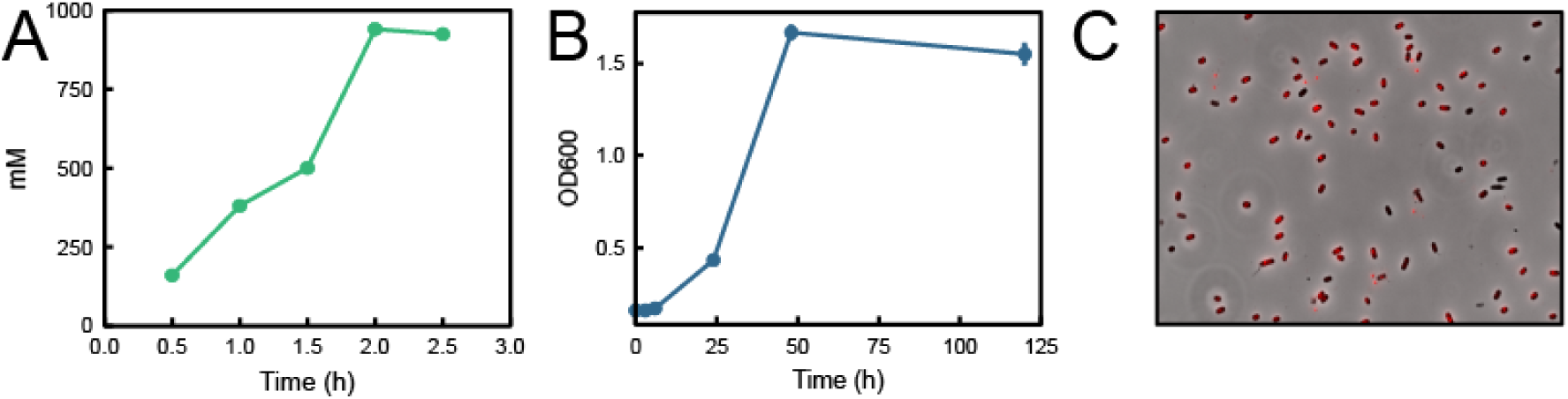
pcPET glycolysis and residual EG upcycling by *P. putida* JM37. (A) pcBHET production during pcPET glycolysis (n = 1). (B) Growth curve of *P. putida* JM37 with residual EG in MC minimal medium (n = 3). (C) Nile Red staining of *P. putida* JM37 growing in the residual EG after 48 h, showing PHA accumulation.

### 3.5. Chemo-biological upcycling of post-consumer PET to PHA

To demonstrate the feasibility of this approach, we tested the pcBHET obtained by glycolysis as carbon source for Con6. As with commercial BHET, 5 mM pcBHET was hydrolysed in 24 h, and MHET was not detected during the experiment (Fig. 5B). In the co-cultures, 68.55 % of TPA and the totality of EG from pcBHET was consumed in 48 h, obtaining 1.36x10^9^ CFU/mL in 48 h in the growth assays. Surprisingly, in this case the *C. testosteroni* 5MH – *P. putida* 3-FP proportion was nearly 50 % - 50 % across the incubation period. Although 9.05 % more TPA was assimilated in these conditions, a 1.55-fold decrease in *C. testosteroni* 5-MH maximum growth was observed compared cultures with commercial BHET, while the growth curve of *P. putida* 3-FP was similar in both cases.

Con6 cultures grown with both BHET and pcBHET were methanolized to quantify PHA production at 24 h. Previous works showed that *C. testosteroni* RW31 accumulates PHB from TPA, while *P. putida* JM37 stores a mixture of mcl-PHA, dominated by the C10 monomer, when growing on EG (Molpeceres-García et al., 2025a; Molpeceres-García et al., 2025b). In this work, Con6 produced predominantly PHB, amounting to 24.7 ± 1.58 % relative to the total cell dry weight of the consortium (%PHB/CDW) in the medium with BHET, and 29.5 ± 0.87 %PHB/CDW when using pcBHET. These values show a notable improvement compared with a previous work in which a mono-culture of *C. testosteroni* RW31 expressing both FAST-PETase and *Is*MHETase accumulated 12.03 %PHB/CDW in 24 h in cultures with 5 mM of BHET (Molpeceres-García et al., 2025). It is known that PHA accumulation can be influenced by nutritional imbalances (Mitra et al., 2022; Montiel-Jarillo et al., 2017). Therefore, these differences could be due to different ratio C/N for both microorganisms in the media, as in Con6 the nitrogen souce is shared between the two populations. Nonetheless, the PHB accumulation mechanisms in *C. testosteroni* RW31 is not well understood yet. These questions should be assessed with further experimental data. Regarding mcl-PHA, the profile showed a predominance of C10 (∼70%), followed by C12 (∼16%) and C8 (∼11%), which slightly differs with that determined for *P. putida* JM37 growing on EG (60% C10; 19% C12; 12% C8; 2% C6) (Molpeceres-García et al., 2025b).

*In vitro* depolymerized PET has been upcycled to a number of valuable compounds, usually employing monocultures (Diao et al., 2023, 2025; Kim et al., 2019; Rebocho et al., 2025). Werner et al. (2021) used glycolysis to depolymerize PET into BHET, which was upcycled to β-ketoadipic acid by a single *P. putida* KT2440 strain modified for EG and TPA assimilation as well as for *Is*PETase and *Is*MHETase secretion. However, glucose was needed as supplementary carbon source to avoid a lag in MHETase activity. In the case of microbial consortia, Bao et al. (2023) designed a *P. putida* KT2440 strain able to assimilate TPA and EG, and compared it with two KT2440 strains, each of them specialized for assimilating either TPA or EG. Employing PET hydrolysate of 75 mM of both TPA and EG, the authors showed that the consortium outperforms the monoculture at upcycling the monomers to mcl-PHA (637 mg/L, 93 % increase) and muconic acid (4.73 g/L, 2.84-fold increase). These results reinforce the concept of applying division of labour within microbial consortia for upcycling.

## 4. Conclusions

This study presents a chemo-biological strategy for the upcycling of post-consumer PET by integrating chemical depolymerization with a synthetic engineered microbial consortium composed of two novel bacterial chassis. Glycolysis effectively converted PET into BHET, overcoming the limitations of *in vivo* degradation. We designed consortia *C. testosteroni* RW31 - *P. putida* JM37, which can respectively assimilate TPA and EG. These bacteria were modified to secrete either a heterologous MHETase or a PETase, allowing the consortium to collaboratively degrade BHET and produce PHA from its monomers. Moreover, various plasmid architectures were tested producing notable differences in population proportions, which could be interesting to design future consortia. We selected a distributed consortium (Con6) based on the growth and production of PHAs, which can be further improved by designing a bioprocess in a bioreactor, optimizing the substrates and inoculum ratios, and controlling the culture conditions. Our work provides proof of concept for the advantages of multi-species microbial consortia in the design of biotechnological strategies, while highlighting the potential of two non-conventional chassis for PET waste valorisation.

## Acknowledgements

This work was supported by the Spanish projects MICODE (PID2020-114210RB-I00 MCIN/AEI), DEMO (MCIN/AEI/“NextGenerationEU”/PRTR, TED2021-130096B-I00) and MOLA (PID2024-162673NB-I00 MCIN/AEI). The authors would also like to thank IBISBA1.0 project (H2020 730976), the SusPlast-CSIC Interdisciplinary Platform, and the LifeHub-CSIC research network for their support. The authors acknowledge the support toward the publication fee by the CSIC Open Access Publication Support Initiative through its Unit of Information Resources for Research (URICI).

## References

1. Aguado, A., Becerra, L., & Martínez, L. (2023). Glycolysis optimisation of different complex PET waste with recovery and reuse of ethylene glycol. Chemical Papers, 77(6), 3293–3303. 10.1007/s11696-023-02704-8

2. Amalia, L., Chang, C.-Y., Wang, S. S.-S., Yeh, Y.-C., & Tsai, S.-L. (2024). Recent advances in the biological depolymerization and upcycling of polyethylene terephthalate. Current Opinion in Biotechnology, 85, 103053. 10.1016/j.copbio.2023.103053

3. Bao, T., Qian, Y., Xin, Y., Collins, J. J., & Lu, T. (2023). Engineering microbial division of labor for plastic upcycling. Nature Communications, 14(1), 1–13. 10.1038/s41467-023-40777-x

4. Barnard, E., Rubio Arias, J. J., & Thielemans, W. (2021). Chemolytic depolymerisation of PET: A review. Green Chemistry, 23(11), 3765–3789. 10.1039/D1GC00887K

5. Barragán, M. J. L., Carmona, M., Zamarro, M. T., Thiele, B., Boll, M., Fuchs, G., García, J. L., & Díaz, E. (2004). The *bzd* Gene Cluster, Coding for Anaerobic Benzoate Catabolism, in Azoarcus sp. Strain CIB. Journal of Bacteriology, 186(17), 5762–5774. 10.1128/JB.186.17.5762-5774.2004

6. Brenner, K., & Arnold, F. H. (2011). Self-Organization, Layered Structure, and Aggregation Enhance Persistence of a Synthetic Biofilm Consortium. PLOS ONE, 6(2), e16791. 10.1371/journal.pone.0016791

7. Bucci, K., Tulio, M., & Rochman, C. M. (2020). What is known and unknown about the effects of plastic pollution: A meta-analysis and systematic review. Ecological Applications, 30(2), e02044. 10.1002/eap.2044

8. Cao, F., Wang, L., Zheng, R., Guo, L., Chen, Y., & Qian, X. (2022). Research and progress of chemical depolymerization of waste PET and high-value application of its depolymerization products. RSC Advances, 12(49), 31564–31576. 10.1039/D2RA06499E

9. Carniel, A., Ferreira Dos Santos, N., Buarque, F. S., Mendes Resende, J. V., Ribeiro, B. D., Marrucho, I. M., Coelho, M. A. Z., & Castro, A.M. (2024). From trash to cash: Current strategies for bio-upcycling of recaptured monomeric building blocks from poly(ethylene terephthalate) (PET) waste. Green Chemistry, 26(10), 5708–5743. 10.1039/D4GC00528G

10. Choi, S. Y., Lee, Y., Yu, H. E., Cho, I. J., Kang, M., & Lee, S. Y. (2023). Sustainable production and degradation of plastics using microbes. Nature Microbiology, 8(12), 2253–2276. 10.1038/s41564-023-01529-1

11. Cook, T. B., Rand, J. M., Nurani, W., Courtney, D. K., Liu, S. A., & Pfleger, B. F. (2018). Genetic tools for reliable gene expression and recombineering in Pseudomonas putida. Journal of Industrial Microbiology & Biotechnology, 45(7), 517–527. 10.1007/s10295-017-2001-5

12. Diao, J., Hu, Y., Tian, Y., Carr, R., & Moon, T. S. (2023a). Upcycling of poly(ethylene terephthalate) to produce high-value bio-products. Cell Reports, 42(1), 111908. 10.1016/j.celrep.2022.111908

13. Diao, J., Hu, Y., Tian, Y., Carr, R., & Moon, T. S. (2023b). Upcycling of poly(ethylene terephthalate) to produce high-value bio-products. Cell Reports, 42(1). 10.1016/j.celrep.2022.111908

14. Diao, J., Tian, Y., Hu, Y., & Moon, T. S. (2025). Producing multiple chemicals through biological upcycling of waste poly(ethylene terephthalate). Trends in Biotechnology, 43(3), 620–646. 10.1016/j.tibtech.2024.10.018

15. Durante-Rodríguez, G., De Lorenzo, V., & Martínez-García, E. (2014). The Standard European Vector Architecture (SEVA) Plasmid Toolkit. In A. Filloux & J.-L. Ramos (Eds), *Pseudomonas Methods and Protocols* (Vol. 1149, pp. 469–478). Springer New York. 10.1007/978-1-4939-0473-0_36

16. Ellis, L. D., Rorrer, N. A., Sullivan, K. P., Otto, M., McGeehan, J. E., Román-Leshkov, Y., Wierckx, N., & Beckham, G. T. (2021). Chemical and biological catalysis for plastics recycling and upcycling. Nature Catalysis, 4(7), 539–556. 10.1038/s41929-021-00648-4

17. Eugenio, E. D. Q., Campisano, I. S. P., De Castro, A. M., Coelho, M. A. Z., & Langone, M. A. P. (2022). Kinetic Modeling of the Post-consumer Poly(Ethylene Terephthalate) Hydrolysis Catalyzed by Cutinase from Humicola insolens. Journal of Polymers and the Environment, 30(4), 1627–1637. 10.1007/s10924-021-02301-4

18. Feng, Y., Lv, S.-W., Zhang, R., Ren, X., Shen, J., & Cong, Y. (2025). From waste to wealth: Glycolysis of PET for high-value resource utilization. Waste Management, 200, 114768. 10.1016/j.wasman.2025.114768

19. Fujita, M., Torigoe, K., Nakada, T., Tsusaki, K., Kubota, M., Sakai, S., & Tsujisaka, Y. (1989). Cloning and nucleotide sequence of the gene (amyP) for maltotetraose-forming amylase from Pseudomonas stutzeri MO-19. Journal of Bacteriology, 171(3), 1333–1339. 10.1128/jb.171.3.1333-1339.1989

20. Gao, R., & Sun, C. (2021). A marine bacterial community capable of degrading poly(ethylene terephthalate) and polyethylene. Journal of Hazardous Materials, 416, 125928. 10.1016/j.jhazmat.2021.125928

21. Hays, S. G., Yan, L. L. W., Silver, P. A., & Ducat, D. C. (2017). Synthetic photosynthetic consortia define interactions leading to robustness and photoproduction. Journal of Biological Engineering, 11(1), 4. 10.1186/s13036-017-0048-5

22. Huang, J., Yan, D., Zhu, Q., Cheng, X., Tang, J., Lu, X., & Xin, J. (2023). Depolymerization of polyethylene terephthalate with glycol under comparatively mild conditions. Polymer Degradation and Stability, 208, 110245. 10.1016/j.polymdegradstab.2022.110245

23. Jia, X., Liu, C., Song, H., Ding, M., Du, J., Ma, Q., & Yuan, Y. (2016). Design, analysis and application of synthetic microbial consortia. Synthetic and Systems Biotechnology, 1(2), 109–117. 10.1016/j.synbio.2016.02.001

24. Johnson, N. W., Valenzuela-Ortega, M., Thorpe, T. W., Era, Y., Kjeldsen, A., Mulholland, K., & Wallace, S. (2025). A biocompatible Lossen rearrangement in Escherichia coli. Nature Chemistry, 17(7), 1020–1026. 10.1038/s41557-025-01845-5

25. Kim, H. T., Kim, J. K., Cha, H. G., Kang, M. J., Lee, H. S., Khang, T. U., Yun, E. J., Lee, D.-H., Song, B. K., Park, S. J., Joo, J. C., & Kim, K. H. (2019). Biological Valorization of Poly(ethylene terephthalate) Monomers for Upcycling Waste PET. ACS Sustainable Chemistry & Engineering, 7(24), 19396–19406. 10.1021/acssuschemeng.9b03908

26. López-Fonseca, R., Duque-Ingunza, I., De Rivas, B., Arnaiz, S., & Gutiérrez-Ortiz, J. I. (2010). Chemical recycling of post-consumer PET wastes by glycolysis in the presence of metal salts. Polymer Degradation and Stability, 95(6), 1022–1028. 10.1016/j.polymdegradstab.2010.03.007

27. Lu, H., Diaz, D. J., Czarnecki, N. J., Zhu, C., Kim, W., Shroff, R., Acosta, D. J., Alexander, B. R., Cole, H. O., Zhang, Y., Lynd, N. A., Ellington, A. D., & Alper, H. S. (2022). Machine learning-aided engineering of hydrolases for PET depolymerization. Nature, 604(7907), 662–667. 10.1038/s41586-022-04599-z

28. Maheswaran, B., Al-Ansari, M., Al-Humaid, L., Sebastin Raj, J., Kim, W., Karmegam, N., & Mohamed Rafi, K. (2023). In vivo degradation of polyethylene terephthalate using microbial isolates from plastic polluted environment. Chemosphere, 310, 136757. 10.1016/j.chemosphere.2022.136757

29. McCarty, N. S., & Ledesma-Amaro, R. (2019). Synthetic Biology Tools to Engineer Microbial Communities for Biotechnology. Trends in Biotechnology, 37(2), 181–197. 10.1016/j.tibtech.2018.11.002

30. Mehta, H., Jimenez, J., Ledesma-Amaro, R., & Stan, G.-B. (2025). Investigating the Potential of Division of Labor in Synthetic Bacterial Communities for the Production of Violacein. ACS Synthetic Biology, 14(7), 2703–2709. 10.1021/acssynbio.5c00120

31. Meyer-Cifuentes, I. E., Werner, J., Jehmlich, N., Will, S. E., Neumann-Schaal, M., & Öztürk, B. (2020). Synergistic biodegradation of aromatic-aliphatic copolyester plastic by a marine microbial consortium. Nature Communications, 11(1), 5790. 10.1038/s41467-020-19583-2

32. Mitra, R., Xu, T., Chen, G., Xiang, H., & Han, J. (2022). An updated overview on the regulatory circuits of polyhydroxyalkanoates synthesis. Microbial Biotechnology, 15(5), 1446–1470. 10.1111/1751-7915.13915

33. Molpeceres-García, F. J., Garcia-Miro, A., Mateos, E., Prieto, A., Sanz, D., Jimenez, J. I., & Barriuso, J. (2025b). Pseudomonas putida JM37 as a novel bacterial chassis for ethylene glycol upcycling (p. 2025.10.07.680640). bioRxiv. 10.1101/2025.10.07.680640

34. Molpeceres-García, F. J., Sanz-Mata, D., García-Miro, A., Prieto, A., & Barriuso, J. (2025a). Towards polyethylene terephthalate valorisation into PHB using an engineered Comamonas testosteroni strain. New Biotechnology, 85, 75–83. 10.1016/j.nbt.2024.12.005

35. Montiel-Jarillo, G., Carrera, J., & Suárez-Ojeda, M. E. (2017). Enrichment of a mixed microbial culture for polyhydroxyalkanoates production: Effect of pH and N and P concentrations. Science of The Total Environment, 583, 300–307. 10.1016/j.scitotenv.2017.01.069

36. Mückschel, B., Simon, O., Klebensberger, J., Graf, N., Rosche, B., Altenbuchner, J., Pfannstiel, J., Huber, A., & Hauer, B. (2012a). Ethylene Glycol Metabolism by Pseudomonas putida. Applied and Environmental Microbiology, 78(24), 8531–8539. 10.1128/AEM.02062-12

37. Mückschel, B., Simon, O., Klebensberger, J., Graf, N., Rosche, B., Altenbuchner, J., Pfannstiel, J., Huber, A., & Hauer, B. (2012b). Ethylene Glycol Metabolism by Pseudomonas putida. Applied and Environmental Microbiology, 78(24), 8531–8539. 10.1128/AEM.02062-12

38. Netzker, T., Flak, M., Krespach, M. K., Stroe, M. C., Weber, J., Schroeckh, V., & Brakhage, A. A. (2018). Microbial interactions trigger the production of antibiotics. Current Opinion in Microbiology, 45, 117–123. 10.1016/j.mib.2018.04.002

39. Nihart, A. J., Garcia, M. A., El Hayek, E., Liu, R., Olewine, M., Kingston, J. D., Castillo, E. F., Gullapalli, R. R., Howard, T., Bleske, B., Scott, J., Gonzalez-Estrella, J., Gross, J. M., Spilde, M., Adolphi, N. L., Gallego, D. F., Jarrell, H. S., Dvorscak, G., Zuluaga-Ruiz, M. E., … Campen, M. J. (2025). Bioaccumulation of microplastics in decedent human brains. Nature Medicine, 31(4), 1114–1119. 10.1038/s41591-024-03453-1

40. OECD. (2022). *Global Plastics Outlook: Policy Scenarios to* 2060. OECD. 10.1787/aa1edf33-en

41. Palm, G. J., Reisky, L., Böttcher, D., Müller, H., Michels, E. A. P., Walczak, M. C., Berndt, L., Weiss, M. S., Bornscheuer, U. T., & Weber, G. (2019). Structure of the plastic-degrading Ideonella sakaiensis MHETase bound to a substrate. Nature Communications, 10(1), 1717. 10.1038/s41467-019-09326-3

42. Pan, L., Khan, W. H., Li, J., Kubota, K., Wang, Q., & Wang, Y. (2025). Metabolic mechanism in biosynthesis of polyhydroxyalkanoate from terephthalic acid by mixed microbial consortium. Chemical Engineering Journal, 515, 163695. 10.1016/j.cej.2025.163695

43. Qi, X., Ma, Y., Chang, H., Li, B., Ding, M., & Yuan, Y. (2021). Evaluation of PET Degradation Using Artificial Microbial Consortia. Frontiers in Microbiology, 12, 778828. 10.3389/fmicb.2021.778828

44. Rebocho, A. T., Torres, C. A. V., Koninckx, H., Stragier, L., Attallah, O. A., Mojicevic, M., Tas, C. E., Fournet, M. B., Reis, M. A., & Freitas, F. (2025). Upcycling depolymerized PET waste into polyhydroxyalkanoates and triacylglycerols by a newly isolated Rhodococcus sp. Strain. Biotechnology for the Environment, 2(1), 5. 10.1186/s44314-025-00019-4

45. Roberts, C., Edwards, S., Vague, M., León-Zayas, R., Scheffer, H., Chan, G., Swartz, N. A., & Mellies, J. L. (2020). Environmental Consortium Containing *Pseudomonas* and *Bacillus* Species Synergistically Degrades Polyethylene Terephthalate Plastic. mSphere, 5(6), e01151–20. 10.1128/mSphere.01151-20

46. Sadler, J. C., & Wallace, S. (2021). Microbial synthesis of vanillin from waste poly(ethylene terephthalate). Green Chemistry, 23(13), 4665–4672. 10.1039/D1GC00931A

47. Seo, H., Kim, S., Son, H. F., Sagong, H.-Y., Joo, S., & Kim, K.-J. (2019). Production of extracellular PETase from Ideonella sakaiensis using sec-dependent signal peptides in E. coli. Biochemical and Biophysical Research Communications, 508(1), 250–255. 10.1016/j.bbrc.2018.11.087

48. Sonnendecker, C., Oeser, J., Richter, P. K., Hille, P., Zhao, Z., Fischer, C., Lippold, H., Blázquez-Sánchez, P., Engelberger, F., Ramírez-Sarmiento, C. A., Oeser, T., Lihanova, Y., Frank, R., Jahnke, H., Billig, S., Abel, B., Sträter, N., Matysik, J., & Zimmermann, W. (2022). Low Carbon Footprint Recycling of Post-Consumer PET Plastic with a Metagenomic Polyester Hydrolase. ChemSusChem, 15(9), e202101062. 10.1002/cssc.202101062

49. Stephens, K., & Bentley, W. E. (2020). Synthetic Biology for Manipulating Quorum Sensing in Microbial Consortia. Trends in Microbiology, 28(8), 633–643. 10.1016/j.tim.2020.03.009

50. Sulaiman, S., Yamato, S., Kanaya, E., Kim, J.-J., Koga, Y., Takano, K., & Kanaya, S. (2012). Isolation of a Novel Cutinase Homolog with Polyethylene Terephthalate-Degrading Activity from Leaf-Branch Compost by Using a Metagenomic Approach. Applied and Environmental Microbiology, 78(5), 1556–1562. 10.1128/AEM.06725-11

51. Thushari, G. G. N., & Senevirathna, J. D. M. (2020). Plastic pollution in the marine environment. Heliyon, 6(8), e04709. 10.1016/j.heliyon.2020.e04709

52. Welsing, G., Wolter, B., Kleinert, G. E. K., Göttsch, F., Besenmatter, W., Xue, R., Mauri, A., Steffens, D., Köbbing, S., Dong, W., Jiang, M., Bornscheuer, U. T., Wei, R., Tiso, T., & Blank, L. M. (2025). Two-step biocatalytic conversion of post-consumer polyethylene terephthalate into value-added products facilitated by genetic and bioprocess engineering. Bioresource Technology, 417, 131837. 10.1016/j.biortech.2024.131837

53. Werner, A. Z., Clare, R., Mand, T. D., Pardo, I., Ramirez, K. J., Haugen, S. J., Bratti, F., Dexter, G. N., Elmore, J. R., Huenemann, J. D., Peabody, G. L., Johnson, C. W., Rorrer, N. A., Salvachúa, D., Guss, A. M., & Beckham, G. T. (2021). Tandem chemical deconstruction and biological upcycling of poly(ethylene terephthalate) to β-ketoadipic acid by Pseudomonas putida KT2440. Metabolic Engineering, 67, 250–261. 10.1016/j.ymben.2021.07.005

54. Wilkes, R. A., Waldbauer, J., Carroll, A., Nieto-Domínguez, M., Parker, D. J., Zhang, L., Guss, A. M., & Aristilde, L. (2023). Complex regulation in a Comamonas platform for diverse aromatic carbon metabolism. Nature Chemical Biology, 19(5), 651–662. 10.1038/s41589-022-01237-7

55. Wright, S. L., & Kelly, F. J. (2017). Plastic and Human Health: A Micro Issue? Environmental Science & Technology, 51(12), 6634–6647. 10.1021/acs.est.7b00423

56. Yao, J., Liu, Y., Gu, Z., Zhang, L., & Guo, Z. (2024). Deconstructing PET: Advances in enzyme engineering for sustainable plastic degradation. Chemical Engineering Journal, 497, 154183. 10.1016/j.cej.2024.154183

57. Yoshida, S., Hiraga, K., Takehana, T., Taniguchi, I., Yamaji, H., Maeda, Y., Toyohara, K., Miyamoto, K., Kimura, Y., & Oda, K. (2016). A bacterium that degrades and assimilates poly(ethylene terephthalate). Science, 351(6278), 1196–1199. 10.1126/science.aad6359

58. Zhang, J., Wang, H., Luo, Z., Yang, Z., Zhang, Z., Wang, P., Li, M., Zhang, Y., Feng, Y., Lu, D., & Zhu, Y. (2023). Computational design of highly efficient thermostable MHET hydrolases and dual enzyme system for PET recycling. Communications Biology, 6(1), 1135. 10.1038/s42003-023-05523-5

59. Zuriani, R., Vigneswari, S., Azizan, M. N. M., Majid, M. I. A., & Amirul, A. A. (2013). A high throughput Nile red fluorescence method for rapid quantification of intracellular bacterial polyhydroxyalkanoates. Biotechnology and Bioprocess Engineering, 18(3), 472–478. 10.1007/s12257-012-0607-z

