## Supplementary Material for "Biochemical Upcycling of PET via Glycolysis and Engineered Microbial Consortia"

**Supplementary Information**

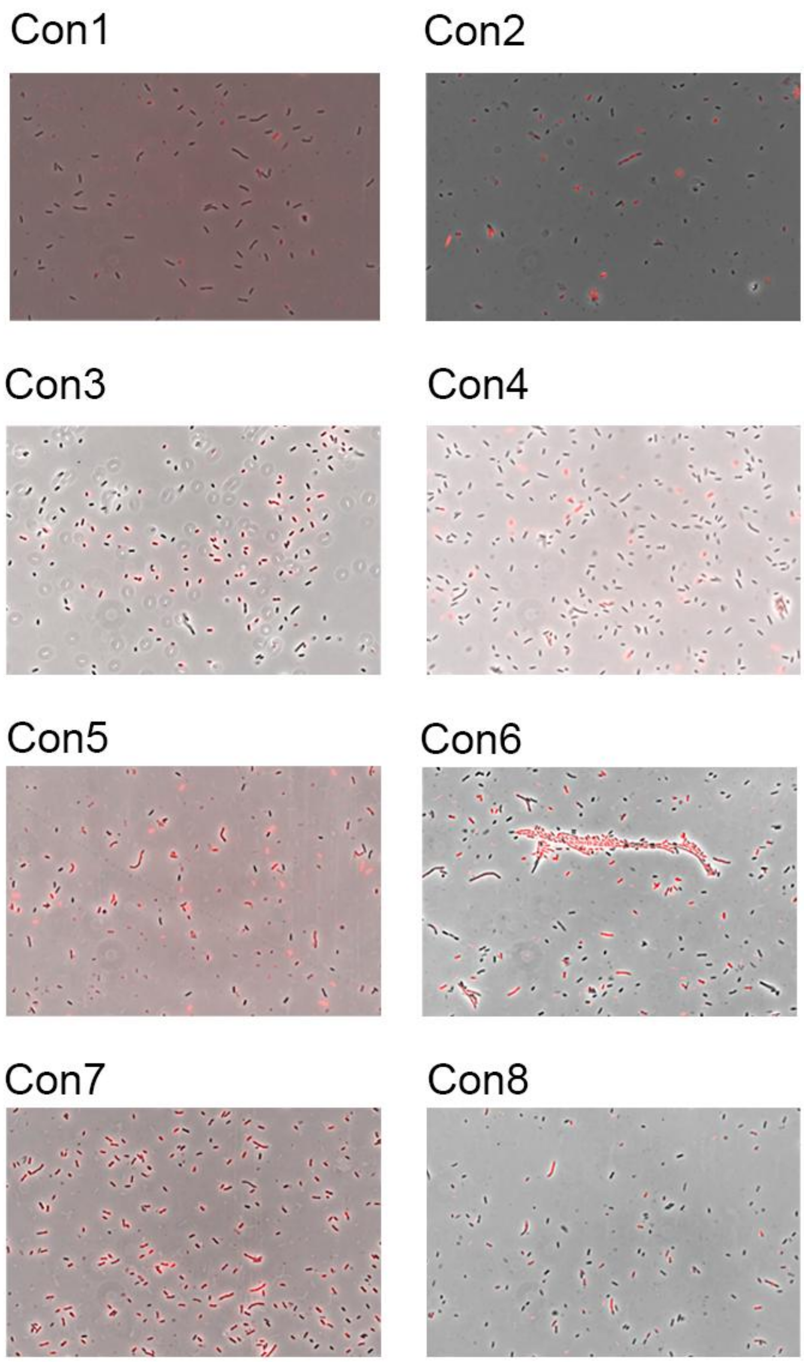


Figure S1. Nile Red staining of the tested consortia at 24 h grown on MC minimal medium with TPA, EG and BHET 5 mM as carbon sources and $NH_4Cl$ 0.1 g/L as nitrogen source (overlayed red and phase contrast channels).
